# Dynamic emperor penguin colonies

**DOI:** 10.64898/2025.12.12.693874

**Authors:** Peter T. Fretwell, Stewart S.R. Jamieson, Grant J. Macdonald

## Abstract

Here we report on the discovery of five emperor penguin breeding locations in West Antarctica, all of which have moved significant distances over the last decade. Three of the colonies could be considered previously undiscovered, while the other two are likely to be relocations due to ice shelf calving events. One of the sites, on the northwest tip of the Stancomb-Wills Ice Tongue has formed or significantly increased in size since 2022, possibly due to immigration from one or both of the two nearest colonies at Halley Bay and Stancomb-Wills colonies. Another colony near Case Island in the Bellinghausen Sea that has recently form is likely due to emigration from other colonies in the region that have suffered recent breeding failures due to early fast ice loss. We discuss the movement of these colonies and the implications for the wider metapopulation and our understanding of emperor penguins’ colony dispersal.

## Introduction

Emperor penguins (Aptenodytes forsteri) are ice obligate seabirds, that breed, forage and moult in the sea ice around the Antarctic continent. They are central place foragers, forming breeding colonies mainly on land-fast sea ice around the continental coastline (Stonehouse 1953). At present there are 66 known breeding sites (Fretwell 2024a), most of which are monitored annually by satellite imagery (LaRue et al 2024). Due to their intrinsic dependence upon sea ice, and based on modelled projections of sea ice change (Diamond et al. 2024, Jenouvrier et al. 2009;2014;2017;2020;2021;2025), their populations are predicted to decline by the end of this century, with a 0-45% chance of extinction depending upon the future climate change scenario (Jenouvrier et al. 2024). Recent changes in sea ice extent and duration have led to a number of catastrophic breeding failures in West and East Antarctica (Fretwell et al. 2023, Wienceke et al. 2024), some of which have been linked to subsequent colony dispersal (Fretwell and Trathan 2019). Several of the monitored colonies have moved location due to changes in the ice coastline (Macdonald et al. 2025, Ancel et al. 2014). This primarily occurs when glaciers and ice shelves close to the colony calve and change the geography and availability of stable fast ice. Furthermore, changes in the duration over which fast ice is present have also been shown to lead to the birds seeking better ice conditions (Fretwell & Trathan 2019). Understanding the adaptability and behaviour of emperor penguins in relation to changing ice conditions is a key parameter in demographic models that assess future extinction risk (Jenouvrier et al. 2017;Jenouvrier et al. 2020). The dynamic nature of emperor penguin colony locations also adds extra complexity to monitoring their populations, as it is sometimes difficult to ascertain whether a colony has moved, or if it is a new population that has not previously been counted (Wienecke 2009, Fretwell 2024a).

Here we describe the discovery of five new colony sites in the Weddell and Bellingshausen Sea, three of which we consider previously undiscovered and two of which are relocations. We track their previous locations and movements and discuss how this information may help understand colony behaviour and future site selection parameters.

## Methods

### The study area

The study area constitutes the region between the eastern Weddell Sea and the western Bellingshausen Sea (Figure 1) between 20° W and 92 ° W. This area was previously known to contain 16 colonies of emperor penguins, several of which have been discovered over the last decade (Fretwell 2024a).

**Figure 1.**
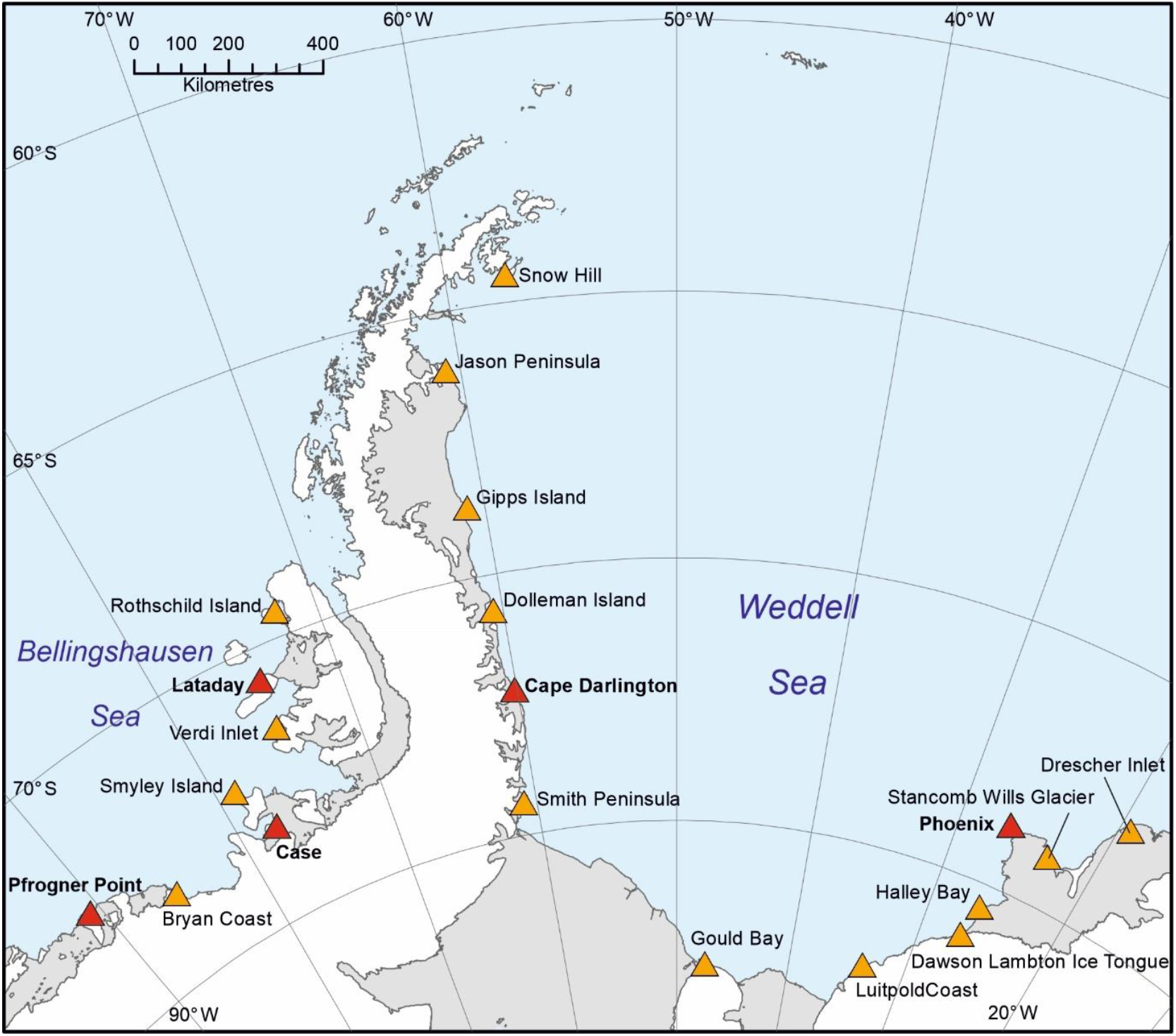
Map of the Weddell to Bellingshausen Sea area of Antarctica. Showing emperor penguin colony locations as of October 2025 Red triangles denote the five colonies discussed in this study, yellow triangles refer to the other know colony locations in the area.

### Satellite imagery

Four Earth Observation satellite platforms were utilized to view emperor penguin colonies. All five of the locations were detected using Sentinel-2 optical imagery. These images are freely available and can be viewed and downloaded on the Copernicus Browser (https://browser.dataspace.copernicus.eu/). They were downloaded as 16-byte georeferenced tiffs into ArcGIS (Esri 2025). The images were viewed using custom parameters, based on a 8/4/3 band combination as detailed in previous work (Fretwell & Trathan 2019, Fretwell 2024b). A list of the images used can be viewed in Supplemental data. To extend the archive further back in time additional Landsat (Landsat 5, 7 and 8) and Aster imagery was viewed in Google Earth Engine using an amended version of the Google Earth Engine Digitisation Tool (GEEDiT) (Lea, 2018).

To confirm the size of the newly discovered Lataday colony, Very High Resolution (VHR) optical satellite imagery was acquired from Vantor (previously MAXAR), using its VHR satellite constellation. Archival imagery from 2024 was available at 50 cm resolution from the WorldView-2 satellite (image ID 30db6b89-7fc2-f000-ba4d-7dc4c0e43562, date 6^th^ October 2024).

To generate an index of springtime abundance, the metric usually used to assess relative colony size (Fretwell et al. 2012, LaRue et al. 2024, Fretwell et al. 2025) a simple binary classifier was constructed in ArcGIS by thresholding the blue band of each image to extract the area of penguins at each site.

For the Phoenix and Case colonies, no archival imagery was available, so only a very rough estimate of five hundred to several thousand can be made from visual assessment of Sentinel-2 imagery.

## Results

We discover emperor penguins in five previous unknown locations in the sector between 20° W and 92° W. Two of these are newly discovered colonies and archival imagery shows that they have been in existence for some time. Case colony has only recently form. The other two; Cape Darlington and Pfrogner Point are previously known breeding aggregations that we show have moved considerable distance due to ice shelf calving events. Below is a brief summary of what we found at each site.

### Lataday colony

A previously undiscovered emperor penguin colony was found at Lataday Island in September 2025. It was first seen in Sentinel-2 imagery located at 74.568° S, 70.670° W from archival imagery taken on October 6^th^ 2024 using the Copernicus Browser (https://browser.dataspace.copernicus.eu/) (Figure 2).

**Figure 2.**
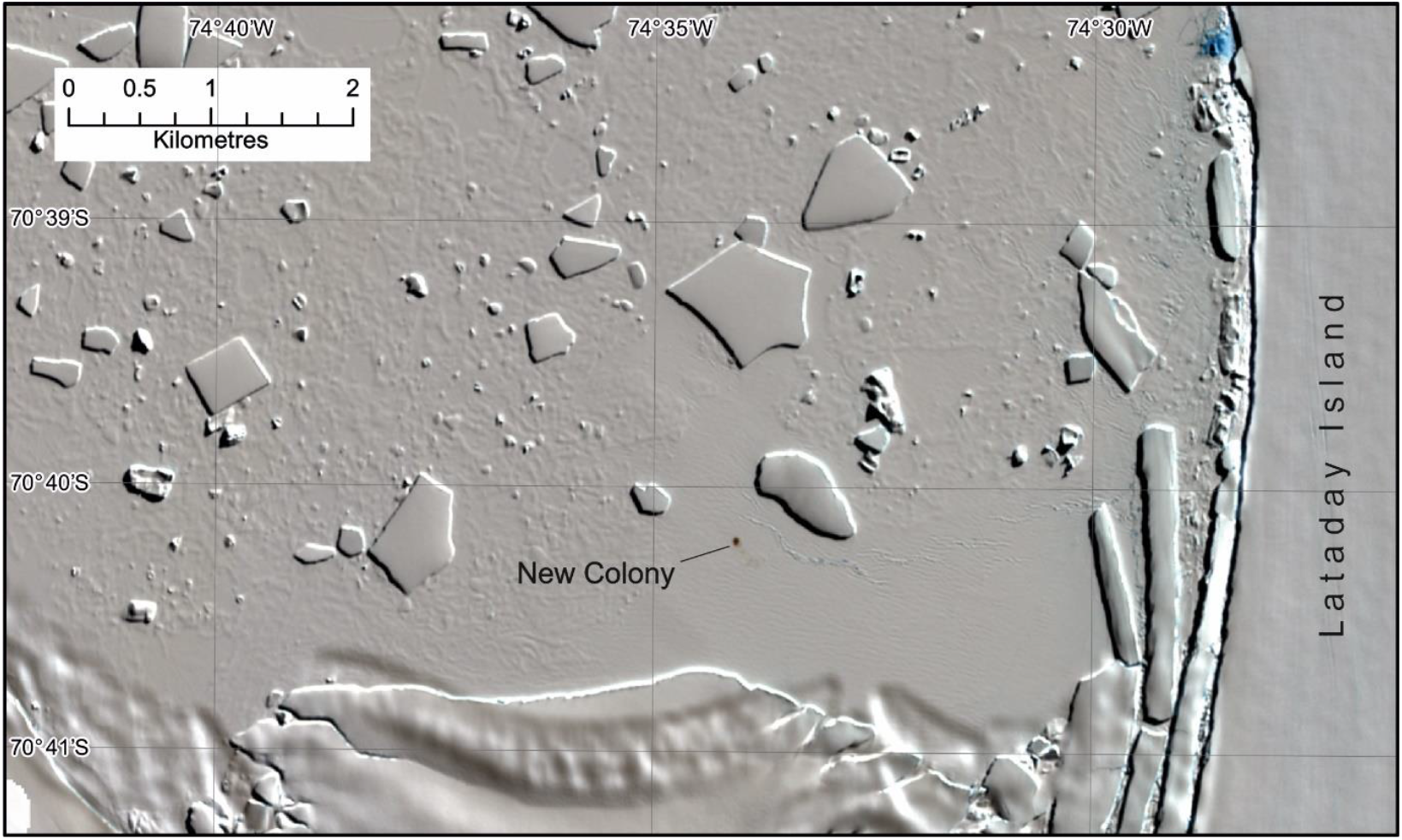
Sentinel-2 image of Lataday colony. This is the original image in which the Lataday colony was discovered at approximately the scale of view, taken on 6/10/2024. Note that a small brown stain, indicative of emperor penguin colonies, is clearly visible on the fast ice less than a kilometre north of a glacier calving from Lataday Island.

In 2024 the colony was located in a deep unnamed embayment on sea ice ~1km north of the coastline of Lataday Island. Lataday Island is an ice-covered landmass 114 km long by 30-50 km wide, lying west of Alexander Island in the Bellinghausen Sea. It is remote and may never have been visited (Hughes et al. 2011). Analysis of archival imagery from Sentinl-2 and Landsat, shows that the colony has been in the bay since 2009, and that before 2009 it was located on sea ice ~40 km west of the present location close to the coast of Lataday Island (Figure 3). The closest known emperor penguin colony is Verdi inlet, ~100 km to the South, although the distance by water would be in excess of 195 km to this site. To the northeast, there is the Rothschild Island colony on the western side of the Antarctic Peninsula. This colony is 150 km northeast as the crow flies, or 180 km by water. The Lataday site is therefore almost equidistant between these two breeding sites. Ancel et al. (2017) proposed that emperor penguin distribution self regulates at intervals around the Antarctic coastline and although the distances between colonies here is less than the average (mean 311 km (Ancel et al. 2017)), it is still some distance from the other sites.

**Figure 3.**
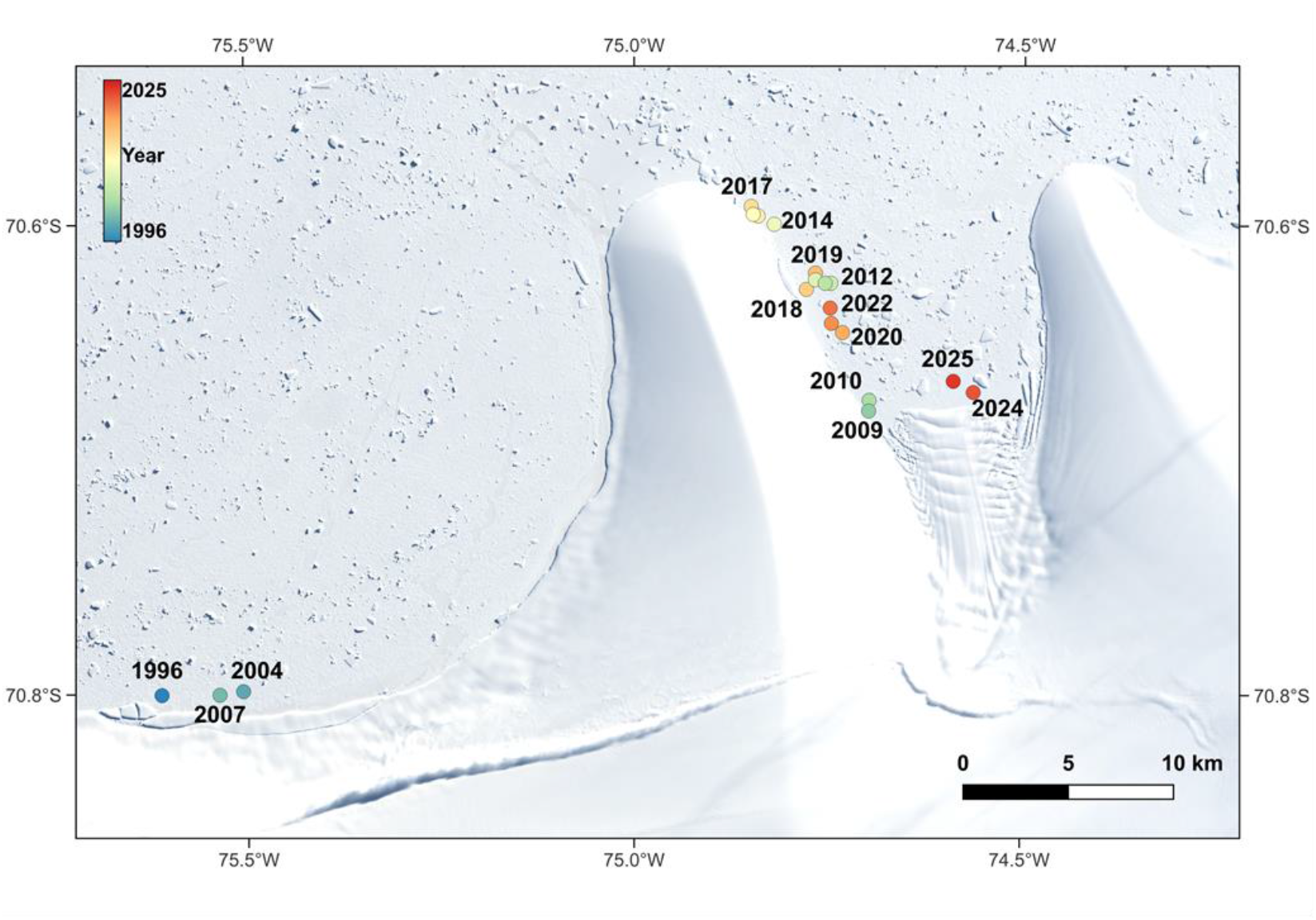
Sentinel-2 satellite image of the northern coast of Lataday island showing the movement of the Lataday emperor penguin colony over time since first visible in Landsat imagery in 1996. The colour of the dot relates to the year the colony was viewed at the location. The colony was first located on the north coast of Lataday Island, near the western end, but in 2009 it had moved into an un-named inlet where it has been since that date.

An assessment of the area of penguins was conducted on VHR optical satellite imagery on 6^th^ October 2024. This used a simple classification method, based on thresholding the panchromatic band to isolate the area of penguins and returned a value of 1426 m^2^. Previous classification analysis indicates density estimates of approximately 1 adult penguin per m^2^ (Fretwell et al. 2012), suggesting over >1400 adult emperor penguins. Springtime estimates do not accurately reflect the total population of a colony, as only a small and variable proportion of the breeding adults are in attendance at this time (Winterl et al. 2024) so this should be seen as a minimum estimate of the true number of penguins using this site.

### Phoenix colony (Stancomb-Wills West)

The first record of this colony was a Bluesky post from a member of the public using the Copernicus browser to search for colonies. The coordinates of this new site were 74.046° S, 26.047° W, near the northwestern corner of the Stancomb-Wills Ice Tongue (Figure 4). This location had not been recorded before and is a considerable distance from other known sites at McDonald Ice Rumples (156 km) and the eastern side of the Stancomb-Wills Ice Tongue (79 km straight line distance, 132 km by sea). Both of these colonies were extant over the time period that this site on the west Stancomb-Wills has been recorded. The colony is located in an embayment in the western side of the ice tongue that is gradually propagating northwards with the ice flow at a rate of approximately 1 km per year. The location has been provisionally named Phoenix colony as it may have grown from the remnants of the now extinct site at Windy Creek (see discussion section).

**Figure 4.**
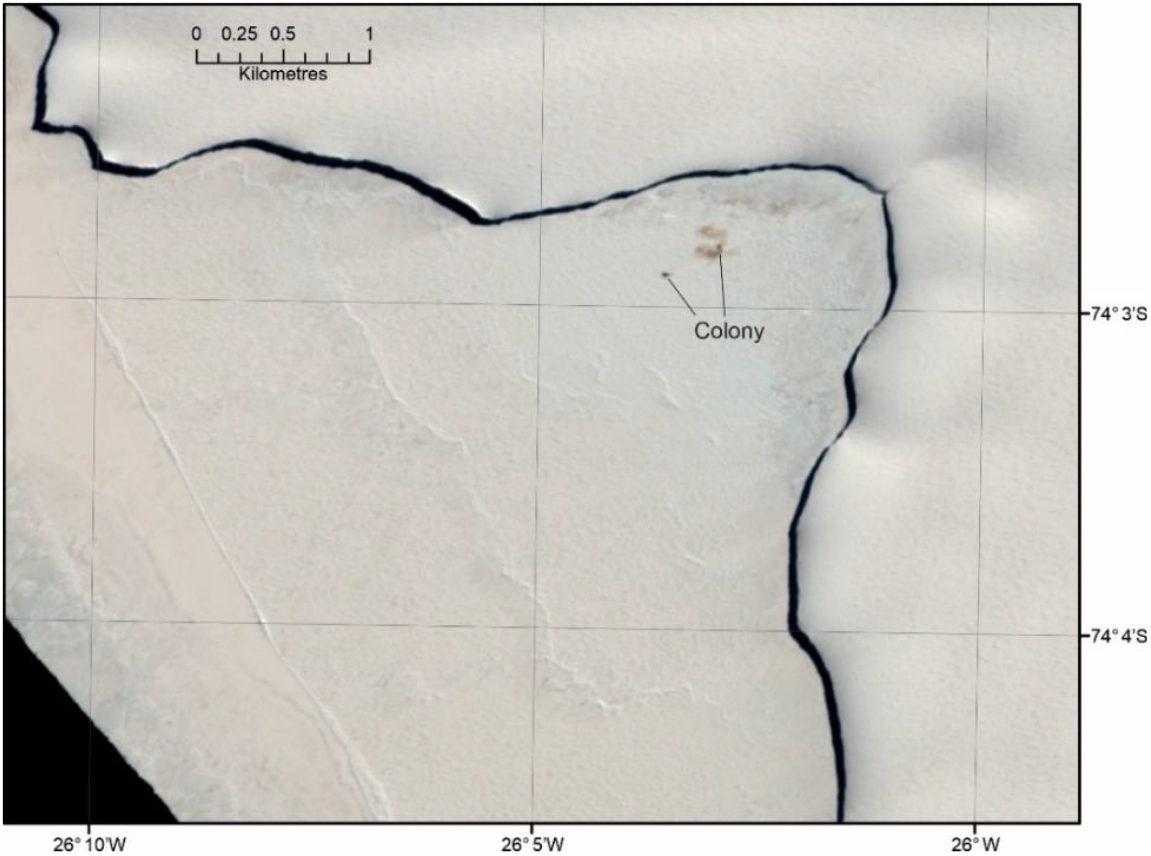
Sentinel-2 image of Phoenix colony (21/10/2025). This image clearly shows the brown guano staining, but also the darker patches that are likely to be the penguins themselves. The un-named embayment on the northwest side of the Stancomb-Wills Ice Tongue has stable multiyear fast ice and is propagating northward at a rate of ~1km per year.

Sentinel-2 archival data, first available in Antarctica in 2016, showed that in October 2016 (6 and 13^th^ November) the staining on the ice, that could have come from the penguins, was very faint and this would have easily been overlooked. In earlier Landsat imagery viewed from the Earth Explorer archive (https://earthexplorer.usgs.gov/) some evidence of a small aggregation of penguins can be seen in 2010 and 2014. Whether this was a breeding site at the time, or just a small aggregation of non-breeders is unknown. With this small level of staining, it would have been impossible to accurately identify it as a colony from these early images without subsequent knowledge. Signs of the colony remained faint between 2017 and 2021, with no staining visible in 2018. But in 2022, the size and colour of the staining had increased in extent and density, indicating that the colony had increase in size significantly. This larger stain remained over the next three years (Figure 5). Springtime counts of individual black dots (likely to be adult emperor penguins, from 30 cm resolution VHR satellite imagery on 13/12/2025 resulted in a count of ~1350 penguins (satellite MAXAR/Vantor Legion1 image ID B110001102DDFA00).

**Figure 5.**
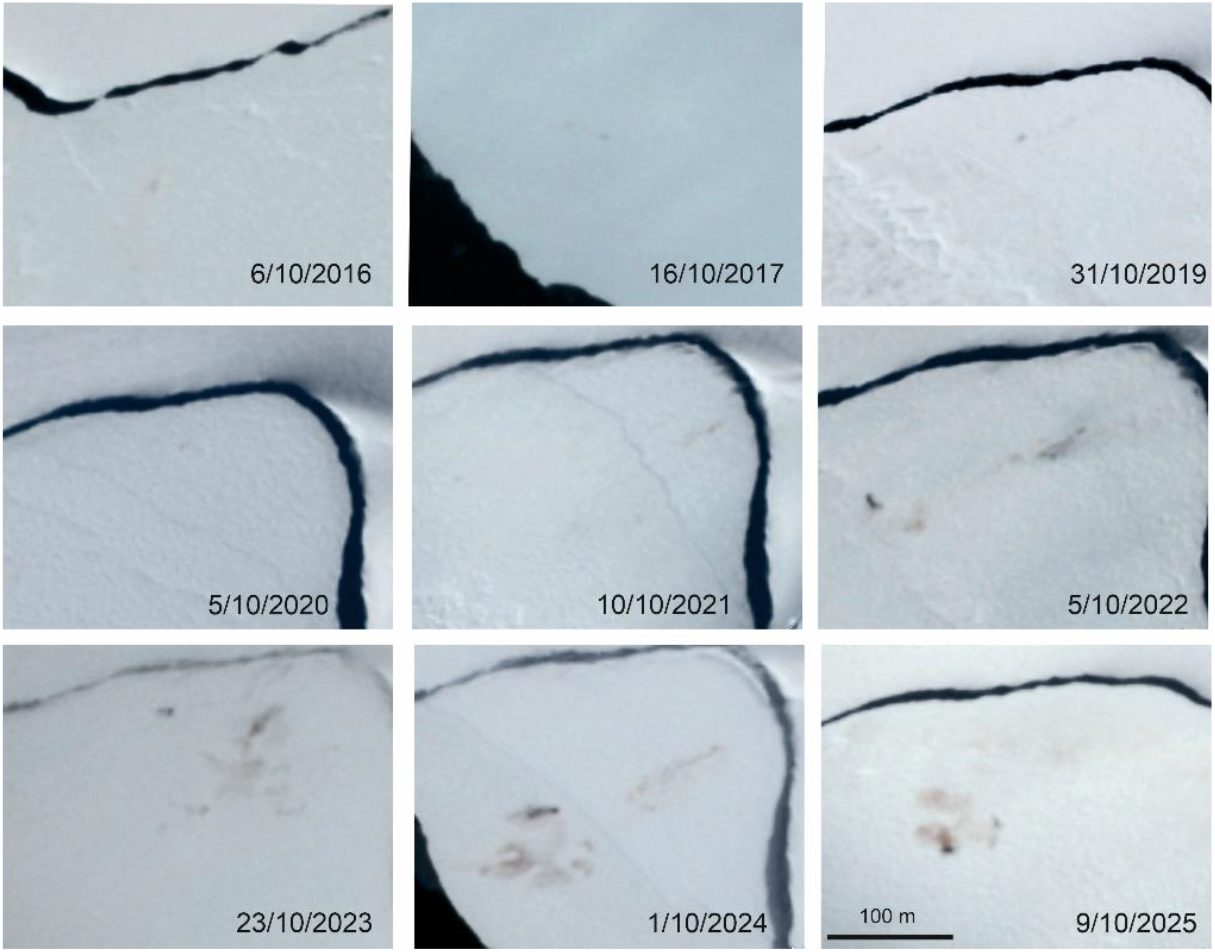
Time series of Phoenix colony from Sentinel-2 imagery. Nine images of the unnamed ice bay on the Stancomb-Wills Ice Tongue, shown at the same scale from 2016 to 2025 (with the exception of 2018 where no staining was visible). Between 2016 to 2021 only very faint signs of penguins were evident, but from 2022 to 2025 a much darker, larger stain was obvious.

**Figure 6.**
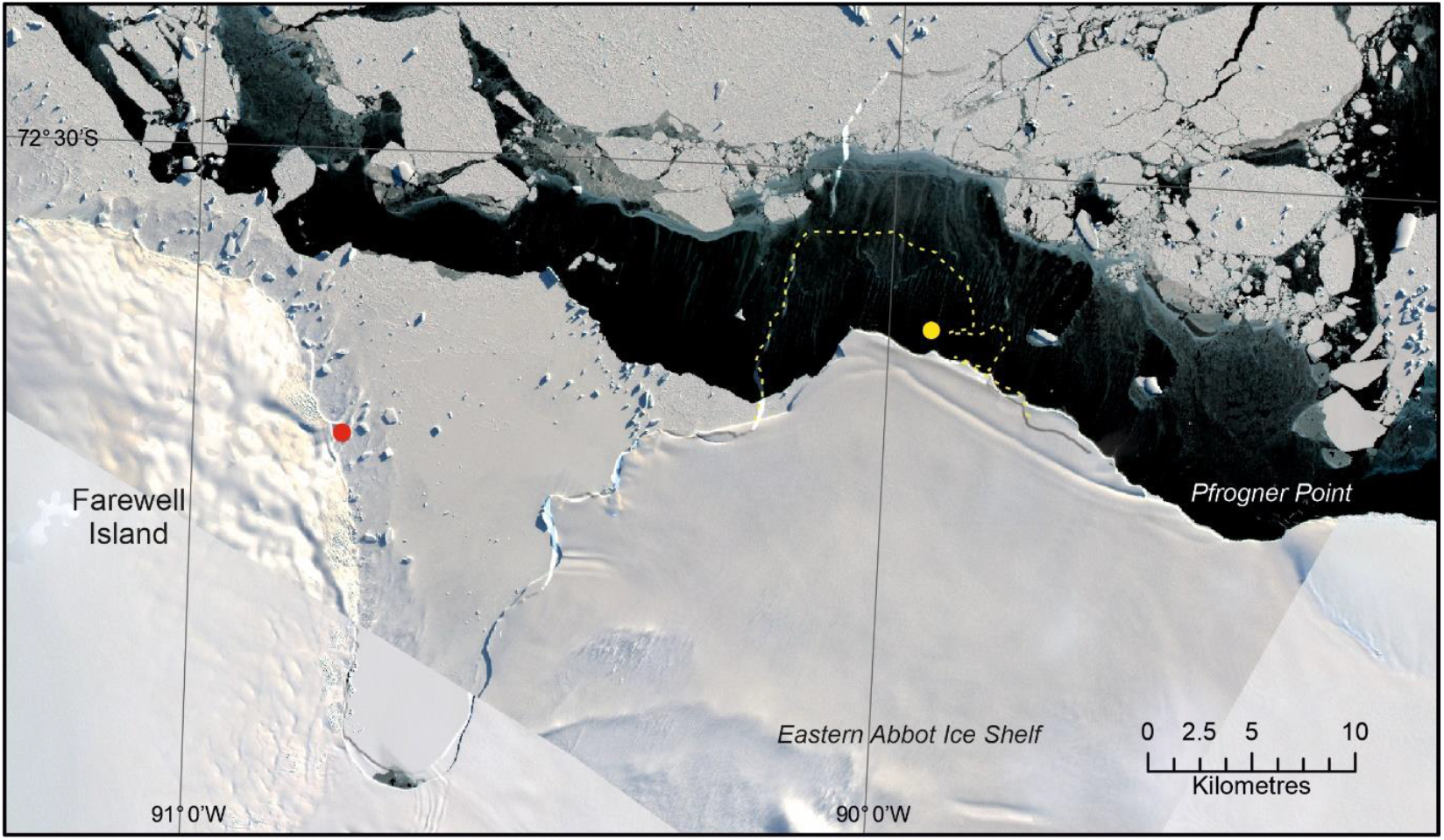
Location of the Pfrogner emperor penguin colony, showing the location in 2018 (yellow dot) and 2025 (red dot), moving from a creek on the eastern Abbot Ice Shelf, to the eastern side of Farewell Island; a movement of 30km. The yellow line shows the extent of the Abbot Ice shelf in 2018, before a significant calving event in 2023.

### Pfrogner Point colony

Pfrogner Point colony was discovered in imagery from 2019 in an ice creek on an ice tongue on the far eastern part of the Abbot Iceshelf at 72.556° S, 89.876°W. Between 2016 and 2022 it was located on the shelf ice at the head of the creek or on the fast ice in the creek itself (Fretwell et al. 2023). Between March 22 and 28^th^ 2023, the ice tongue on which the colony was located calved along the line of the ice creek, taking away the secure breeding location and altering the fast ice regime in the area so that no suitable breeding habitat remained. In late 2023 Sentinel-2 imagery was used to search areas of fast ice in the region for any sign of the re-formation of the colony. October 2023 there were tentative signs of the re-formation of the colony in an unnamed bay 30 km west of the previous location at 72.599° S, 90.822° W near the eastern side of Farewell Island. At this time VHR satellite imagery of the site showed only a few tens of birds (October 2023). In 2024 the colony had moved 3.2 km southwards and the staining on the ice was more obvious. In 2025 the colony remained in this position at 72.624°S, 90.784°W, with more extensive staining. No VHR satellite imagery of this new site has yet been acquired.

### Case Colony

A colony was found on the western side of Case Island in Carroll Inlet, part of the Bellingshausen Sea (-73.164 S, -77.820 W). Like other colonies it was identified using Sentinel-2 imagery, initially from imagery in November and December 2025. Checks of archival imagery show that this colony only formed in 2023, when it was small and indistinct, but it has become more obvious in 2024 and 2025. It is located in a deep embayment and is 105 km (144km by sea) south of the nearest other colony at Smyley Island (Figure 7). Since 2022 Smyley Island and other colonies in the Bellingshausen Sea region have suffered breeding failures from reducing fast ice duration (Fretwell et al. 2023, Fretwell 2024b), so it seems likely that this new colony may be emigrants from those colonies who have prospected for more stable ice. The inner part of Carroll Inlet is one of the few places in the Bellinghausen Sea which still retains stable fast ice for much of the year. However, distance from the colony to the ice edge early in the season can be over 50 km.

**Figure 7.**
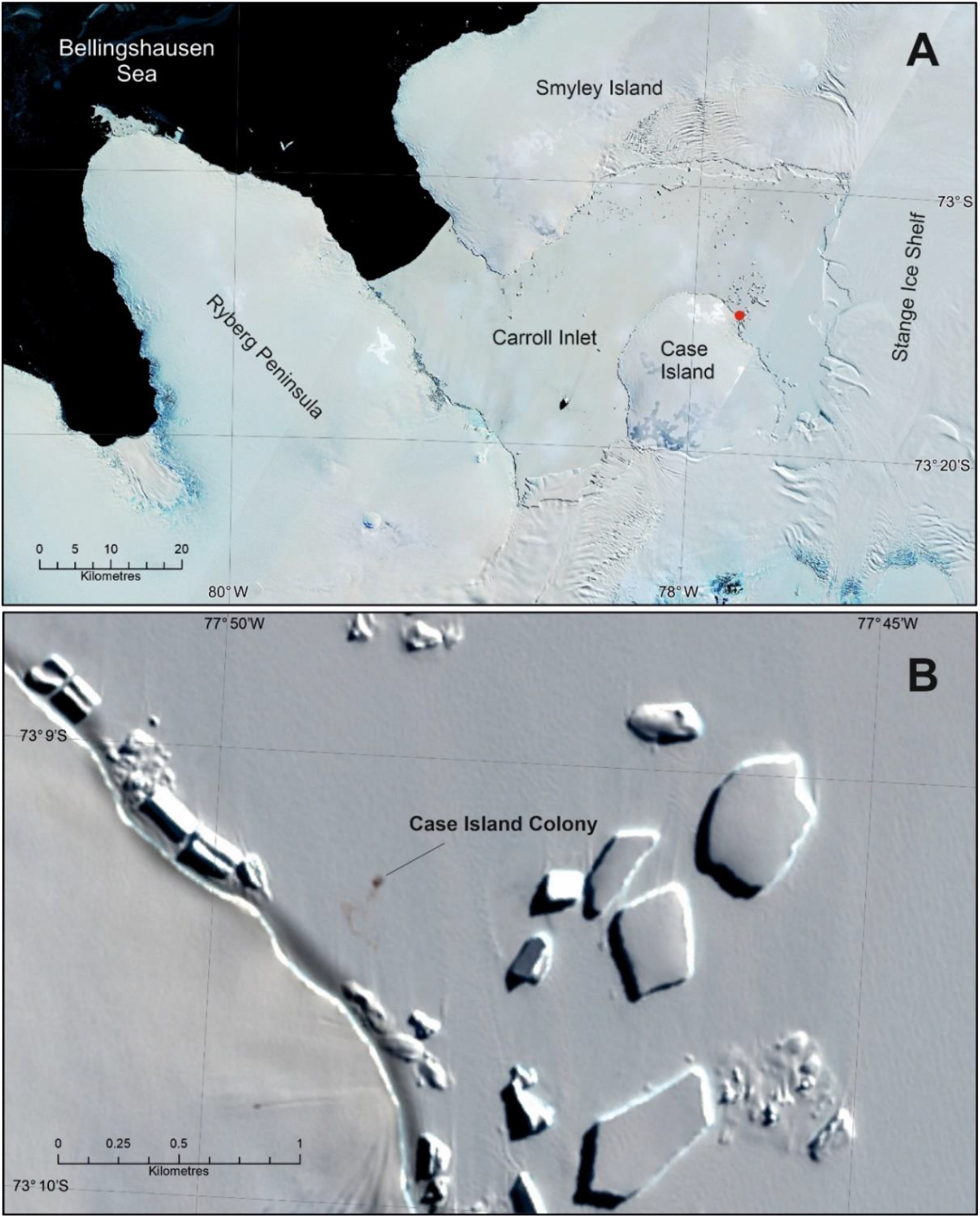
the location of Case colony. A new colony was found near Case Island in Carroll inlet in the Bellingshausen Sea (7A). This colony was obvious in Sentinel-2 imagery from 2025 as a dark patch on the ice with an indicative brown guano stain (7B). Imagery Sentinel-2, 7A from 6 and 10^th^ Oct 2024, 7B from 6^th^ October 2024.

### Cape Darlington colony

The colony was first found in imagery from 2019 on the iceshelf near Cape Darlington on the glacial ice tongue flowing from Hilton Inlet at 71.852° S, 60.125 W (Fretwell and Trathan 2020) (Figure 8a). It remained on the ice shelf until 2021, when it relocated onto the fast ice, just beneath the original 2019 location. However, after early sea ice break-up in 2021, it relocated 9 km south, back onto the ice shelf. In 2022 it moved a further 2.5 km south, back onto the fast ice, where it was clearly visible in September and October. On 28^th^ November that year, before chick fledging, the ice shelf behind the colony broke off, removing the fast ice and changing the local geography considerably. In 2023 the colony had located 45 km south of the 2019 position, to a new location on the fast ice in a small inlet south of Butler Island at 72.260° S, 60.165°W. It has remained close to this location since and is now obvious in satellite imagery (Figure 8b). The colony remains small. In VHR satellite imagery from 2023 (Figure 8c) the emperor penguins are well spaced and the image resolution good enough to count individuals. This gives a springtime estimate of >300 adults. Sentinel-2 imagery in 2023 shows that the colony was relatively small and subsequent imagery in 2024 and 2025 indicates that numbers may have increased since then.

**Figure 8.**
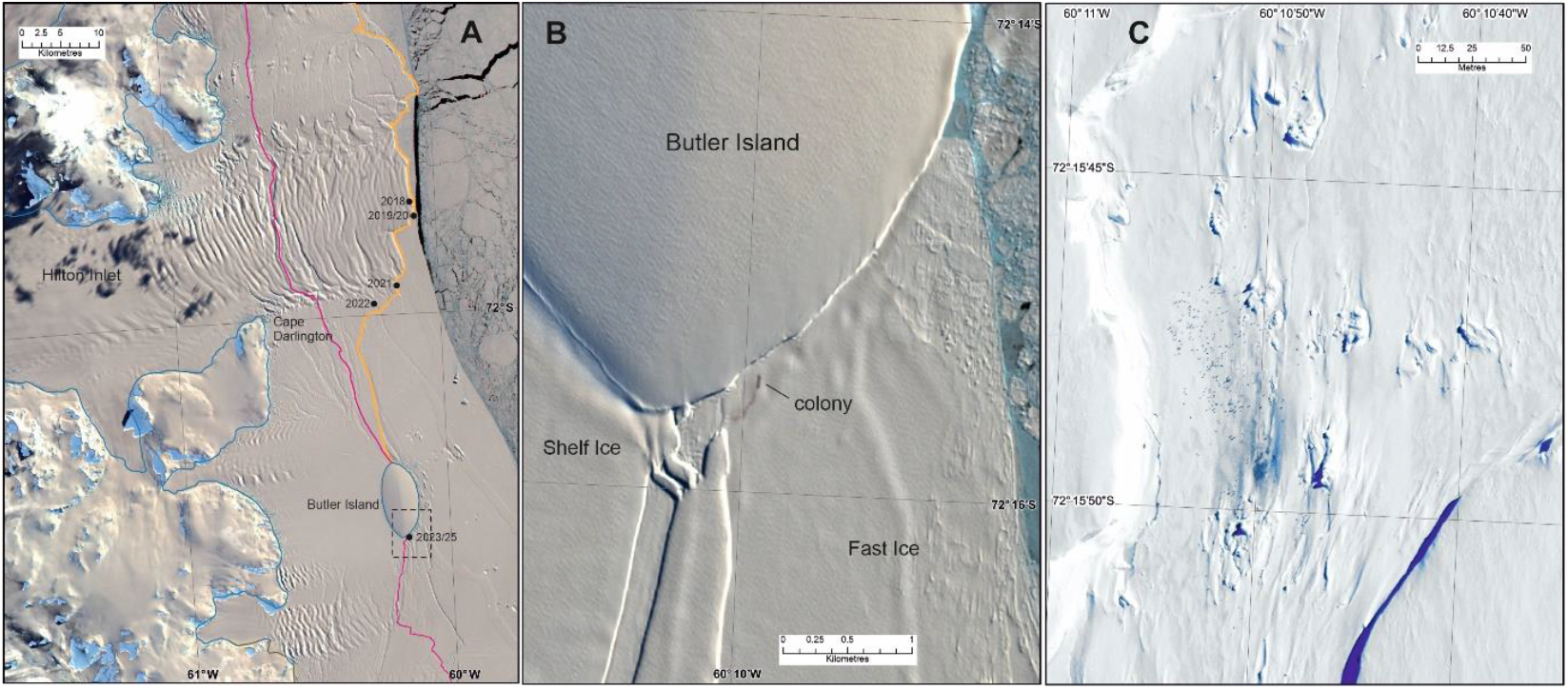
Movements of the Cape Darlington emperor penguin colony. A; overview map showing the southward migration of the colony since 2018. Yellow line denotes the pre-2023 coastline, red line is the post 2023 coastline after the calving event. Blue line denotes the grounding line. The dashed black rectangle denotes the extent of map B (coastline polygons are taken from the Antarctic Digital Database 2025 https://www.bas.ac.uk/project/add/. B:Sentinel-2 image from 17^th^ September 2025 showing the present location of the colony at the southern end of Butler Island. C: Detailed zoom-in of the colony location in 2023 near Butler Island showing dispersed animals that can be counted individually. Imagery from WorldView-3, Image ID 104001008E6D1E00 resolution 0.4m GSD WV3 date 19/11/2023.

## Discussion

### General movements

Movement of emperor penguins is a response to the changing nature of the ice landscape which they inhabit. Unlike species that breed on land, the ever-moving geography of the sea ice and ice sheet coastline ensures that this species has to react to changes in regional geography, changing its location as and when the ice dictates. Seventy percent of the Antarctic coastline is comprised of ice shelves and glaciers and colonies associated with these environments, plus the ones that depend on fast ice that is pinned by grounded icebergs, “have to” relocate occasionally when the surrounding geography changes. At present these criteria account for 40 of the (now) 69 known breeding sites (58%). At present it is unclear what impact movement may have on colony numbers. If an ice shelf or glacier calves during the breeding season, then the colony could lose that year’s cohort of chicks, but there may longer consequences to the metapopulation if the colony, or some of the colony, takes several years to find a new suitable breeding area. The examples shown here, from Cape Darlington, Pfrogner Point and Phoenix colonies suggests that relocated colonies often takes two-to-three years to regain their original size after relocation.

### Movements due to ice shelf calving

The colonies examined in this paper show the dynamic nature of emperor penguin colonies. The movement of Pfrogner and Cape Darlington colonies is easiest to explain. Their movement is mainly due to cyclical ice shelf calving events, part of the natural cycle of ice shelf extension and break-off, although we note that both calving events took place in 2023 in a year of record low sea ice. Low sea ice can lead to increased calving (Liu et al 2015, Teder et al 2025), so climate related factors cannot be completely ruled out as the overarching driver for colony disruption and movement. The Cape Darlington colony has moved progressively south seeking more constant ice conditions, while the Lataday colony has moved eastward into a deep embayment where the fast ice is more stable. This migration may be linked to long-term declining fast ice in the Bellingshausen Sea area (Abram et al. 2010).

### Phoenix colony and Halley Bay

The sudden growth of the Phoenix colony in 2022 cannot be a result of recruitment and can only be explained by immigration from other sites. The closest comparable example is the sudden growth of the Dawson Lambton colony after the collapse of the Halley Bay breeding site between 2016 and 2018 (Fretwell and Trathan 2019), and it is possible that the sudden growth of the Phoenix colony is a direct result of the Halley Bay collapse. Before the Halley collapse the Dawson Lambton colony already had a population of several thousand. Interestingly, prior to the immigration of the Halley penguins to Dawson Lambton it had one colony centre, but after 2018 it formed two distinct centres that were originally 1.3 km apart and these two distinct colonies have continued to slowly drift apart (in 2025 they were 2.5 km apart), a characteristics which seems to be unique to this site. From this observation we have to ask the question; if the sudden growth of Dawson Lambton was from Halley Bay immigrants, did they integrate with the original penguins at Dawson Lambton, or did they form their own sub-colony?

Besides the emigration to Dawson Lambton, after the documented collapse of the colony Halley Bay in 2016, in 2018 a much smaller colony of just a few hundred birds re-formed close to the original “*Windy Creek*” colony location (Fretwell and Trathan 2019). This site persisted over the next few years and was extant in October 2022, when the Phoenix colony appeared (Figure 9). During these years this small Brunt Ice Shelf colony suffered fast-ice loss before the December fledging period every year, so it is likely that few, if any chicks fledged (Fretwell and Trathan, 2019). After the calving of the Brunt Ice Shelf in 2023 the Halley colony moved 32 km east to the eastern side of the McDonald Ice Rumples (from 75.494°S, 27.373°W to 75.453°S, 26.239°W), where it is still clearly visible in Sentinel-2 imagery. Although no population counts have been possible, the size of the colony from Sentinel-2 imagery appears to be similar to the aggregation on the Brunt Ice shelf between 2018 and 2022. It does however remain intriguing that the Phoenix colony appeared the year before the Brunt Ice Shelf calved.

**Figure 9.**
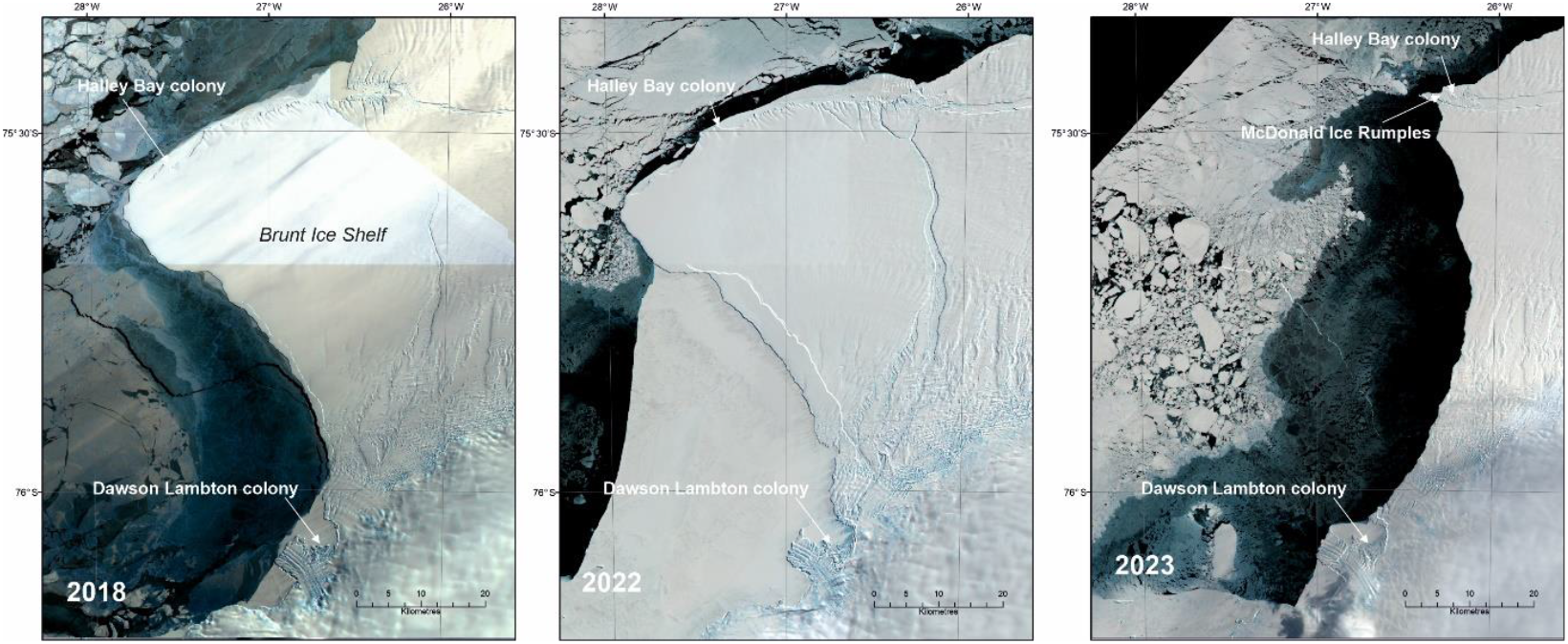
Movement of the Halley Bay emperor penguin colony after 2016. By 2018, most of the emperor penguins from the original Halley Bay site had moved to Dawson Lambton colony, but the colony at Halley Bay reformed on a much smaller scale in 2018. It remained in a similar position, with just a few hundred (from the original 22,000) birds until 2023, when the Brunt Ice Shelf calved and the small colony was forced to relocate 32 km east near the McDonald Ice Rumples. Sentinel-2 images from the Copernicus Browser.

An alternate explanation for the sudden growth of the Phoenix colony is that the majority of the penguins moved from the Stancomb-Wills colony to the east of the ice shelf. Population estimates show that this colony has been decreasing in size since satellite monitoring began in 2009 (LaRue et al. 2024, Fretwell et al. 2025). This could be due to the propagation of the ice shelf, which has increased local sea ice extent and rendered the colony site less desirable because travel to and from the colony from the ice edge takes longer. A similar repositioning occurred in the Lazarev Ice Shelf colony possibly also due to ice tongue propagation (Fretwell 2024a). However, most of the observed population decrease seen in satellite data happened between 2009-2016 and there has been minimal population decline since 2017 (Fretwell et al. 2025 Supplemental 6), which is the period over which we observe the sudden growth of the Phoenix colony. Overall, no corresponding observations in other colonies fully explain the sudden appearance of the Phoenix colony, but its appearance emphasizes the complex nature of regional emperor penguin metapopulation interactions (LaRue et al. 2015).

### Dispersal and emigration

Colony movement and dispersal is expected to become more common as climate change forces emigration of adult penguins from sites that are no longer tenable (Jenouvrier et al. 2017). How emperor penguins adapt and disperse is one of the critical parameters in ongoing attempts to model future emperor penguin population trends. The movements seen in this paper, show that emperor penguins disperse in several ways; moving to existing sites, remaining close to sub-optimal sites or forming new sites. Colonies sometimes move “en-mass”, but may also split (as seen at Halley, SANAE, and Mertz; Fretwell and Trathan, 2019; Macdonald et al., 2025), so the response to environmental stressors can be complex and multifold. It seems however that in most cases emperor penguins make informed choices, seeking out stable ice (as in the example of Case colony), sometimes many kilometres away, and do not move randomly to unsuitable ice conditions (one of the possible scenarios of previous models).

With decreasing sea ice, warming oceans, accelerated ice calving and increased storminess impacting fast ice extent and duration (Fretwell et al. 2023), the ability to relocate to better ice conditions will be critical for the survival of emperor penguins. Understanding the ability and propensity of the dispersal of emperor penguins from sub-optimal breeding sites to find sites with “better ice”, and the potential impact on the breeding success of metapopulations is a key component in demographic models (Jenouvrier et al. 2020). This paper shows that movement of colonies tens to hundreds of kilometres is possible and happens on a routine basis, but movement over larger distances is as yet unrecorded.

## Conclusion

We find five new site locations for emperor penguin colonies, three of which are previously undiscovered. We discuss the movements of colonies and the potential impact of moving on metapopulations, including the complex history of the Halley Bay/McDonald Ice Rumples colony. The movement of colonies can be caused by a number of factors including decreasing sea ice extent, reducing fast ice duration and increased iceberg calving. Sudden forced movement can lead to breeding failure at the time of the event and reduced breeding success in subsequent years. With climate change it is likely that these drivers will intensify in future years, leading to more movement of colonies, with associated breeding failures, which, over time, will impact the overall population. Studying how emperor penguins react and adapt to these changing conditions will be crucial to understand future population trajectories.

The dynamic nature of colonies and the unexpected discovery of new sites, also poses potential challenges for ongoing efforts to monitor the species. This is especially problematic if colonies can suddenly form 100s of km from existing sites. This suggests that if we are to understand emperor penguin populations, the whole coastline of the continent must be monitored each year to ensure no colonies are missed.

## Data statement

Shapefile with 2025 colony locations available from Supplemental table 1 Sentinel-2 imagery available from Copernicus Browser. Images used in this analysis Supplemental table 2

## Acknowledgements

We would like to acknowledge the funding for satellite imagery and analysis time from WWF UK (GB095701), project NEB2181 “Understanding emperor penguin populations in the Weddell Sea and Antarctic Peninsula” and previous WWF funding over the 15-year period. We also acknowledge Natural Environment Research Council (NERC) UK NE/Y00115X/1 awarded to SSRJ and PTF

We would like to sincerely thank Rafał Wereszczyński for the original discovery of the new colony of the west side of Stancomb-Wills Ice tongue and the posting of it and its coordinates on Bluesky.

This paper is linked the Southern Ocean Observing System’s Censusing Animal Populations from Space (CAPS) Special Action Group.

